# Persistence of quantal synaptic vesicle recycling following dynamin depletion

**DOI:** 10.1101/2020.06.12.147975

**Authors:** Olusoji A.T. Afuwape, Natali L. Chanaday, Merve Kasap, Lisa M. Monteggia, Ege T. Kavalali

## Abstract

Dynamins are GTPases required for pinching vesicles off the plasma membrane once a critical curvature is reached during endocytosis. Here, we probed dynamin function in central synapses by depleting all three dynamin isoforms in postnatal hippocampal neurons. We found a decrease in the propensity of evoked neurotransmission as well as a reduction in synaptic vesicle numbers. Using the fluorescent reporter vGluT1-pHluorin, we observed that compensatory endocytosis after 20 Hz stimulation was arrested in ~40% of presynaptic boutons, while remaining synapses showed only a modest effect suggesting the existence of a dynamin-independent endocytic pathway in central synapses. Surprisingly, we found that the retrieval of single synaptic vesicles, after either evoked or spontaneous fusion, was largely impervious to disruption of dynamins. Overall, our results suggest that classical dynamin-dependent endocytosis is not essential for retrieval of synaptic vesicle proteins after quantal single synaptic vesicle fusion.

## Introduction

The robustness of synaptic transmission relies on the ability of retrieving, reforming and refilling synaptic vesicles after their fusion in a reliable and swift manner (Chanaday and Kavalali, 2017). Although the molecular machinery responsible for regulated secretion and endocytosis is conserved among species and cell types, some components have evolved to be more specific for neurons (Saheki and De Camilli, 2012; Soykan et al., 2016). During endocytosis, a critical step involves scission of the nascent vesicle from the plasma membrane. The observation of depleted synaptic vesicles and arrested budding endosomes in the shibire mutant fly (Koenig and Ikeda, 1989) led to the discovery of the GTPase dynamin as the essential protein catalyzing the fission of membranes (Antonny et al., 2016; Ferguson and De Camilli, 2012).

While *Drosophila* has a single gene for dynamin, mammalians have three dynamin isoforms, all of which are expressed in the nervous system and are present at the synapse (Ferguson and De Camilli, 2012). Defects in synaptic vesicle recycling during high frequency stimulation have been described after deletion of the most abundant dynamin isoforms — dynamin 1 and 3 — in the nervous system; however, synaptic vesicle endocytosis persisted partly due to the more ubiquitously expressed isoform dynamin 2 (Ferguson et al., 2007; Raimondi et al., 2011). In superior cervical ganglion neurons, knock down of dynamin 2 revealed defects in rapid recovery of the readily releasable pool of synaptic vesicles after high frequency stimulation (Tanifuji et al., 2013). A similar finding was reported in chromaffin cells where after prolonged stimulation, a dynamin 2 dependent endocytosis mechanism was activated to retrieve vesicles (Artalejo et al., 2002). A function for dynamin 2 at the synapse is also supported by the finding that Ca^2+^ influx in neurons inhibits dynamin mediated endocytosis at the active zone and has been shown to reduce specifically dynamin 2 GTPase activity in HeLa cells (Cousin and Robinson, 2000). Taken together, all the different isoforms of dynamin have been functionally implicated in synaptic vesicle endocytosis with varying degrees of importance but with substantial overlap among them.

Assessing the role of all dynamin isoforms has been challenging due to the non-viability of dynamin triple knock out (TKO) animals. Instead, researchers have turned to the use of dynamin inhibitors such as dynasore and dyngo-4a which allowed elucidating the role of dynamin in multiple forms of endocytosis that reportedly operate at the synapse (Linares-Clemente et al., 2015; McCluskey et al., 2013; Watanabe et al., 2013). However, these dynamin inhibitors have been shown to also inhibit endocytosis in dynamin TKO fibroblasts suggesting that prior and future work with such inhibitors should be interpreted with caution (Park et al., 2013). Moreover, off-target effects on other presynaptic pathways have been described (Douthitt et al., 2011).

In this study, we show that synaptic vesicle recycling and neurotransmission in cultured hippocampal neurons are unaffected by postnatal deletion of dynamin 2 indicating that this dynamin isoform is not essential for synaptic function. Moreover, we were able to deplete all three dynamin isoforms in cultures and found a decrease in the number of synaptic vesicles and number of docked vesicles within the synapse suggesting a role for dynamins in synaptic vesicle pool maintenance. More importantly, while the previously reported arrest in endocytosis was observed after 20 Hz stimulation using the fluorescent reporter vGluT1-pHluorin, we show that it occurs only in a fraction of all presynaptic boutons (~40%). The rest of synapses show only an intermediate effect or the complete absence of it revealing the presence of dynamin-independent endocytosis in central synapses. In line with this notion, we found that this dynamin-independent recycling of individual synaptic vesicles, after either evoked or spontaneous fusion, was largely impervious to disruption of actin, Arp2/3 and the dynamin-related protein (DRP) function. Our results suggest that retrieval of synaptic vesicle proteins does not require classical dynamin-dependent endocytic pathways after the release of single synaptic vesicles.

## Results

### Dynamin 2 is not essential for synaptic vesicle recycling and neurotransmission

Dynamin 2 catalyzes the scission of budding endosomes for various types of endocytosis apart from clathrin-mediated endocytosis (Cao et al., 2007; Liu et al., 2008; Schlunck et al., 2004) and it was reported to play a role in the exocytosis-endocytosis coupling of vesicles in mouse pancreatic β-cells (Min et al., 2007). In neurons, dynamin 2 has been reported to partially compensate for dynamin 1 knock out (KO) phenotype (Raimondi et al., 2011). Taken together, these findings hint at a potential role for dynamin 2 in synaptic vesicle recycling. Complete loss of dynamin 2 has proven to be too strenuous on developing tissue and as such, a complete dynamin 2 KO mouse is non-viable in utero (Ferguson and De Camilli, 2012). Here, we successfully knocked out dynamin 2 postnatally in dissociated hippocampal neuron cultures using dynamin 2 floxed (dnm2f/f) mice and lentiviral delivery of the Cre recombinase. We confirmed dynamin 2 conditional knock out (Dnm2 cKO) by western blot analysis (Figure 1A). At 18 DIV, dynamin 2 levels were reduced by (92.5±1.5)% in neurons expressing the Cre recombinase (Dnm2 cKO) compared to control neurons (Figure 1A). Maximal depletion of dynamin 2 was observed after 17 DIV and thus all subsequent experiments were conducted at 17 DIV or later.

**Figure 1.**
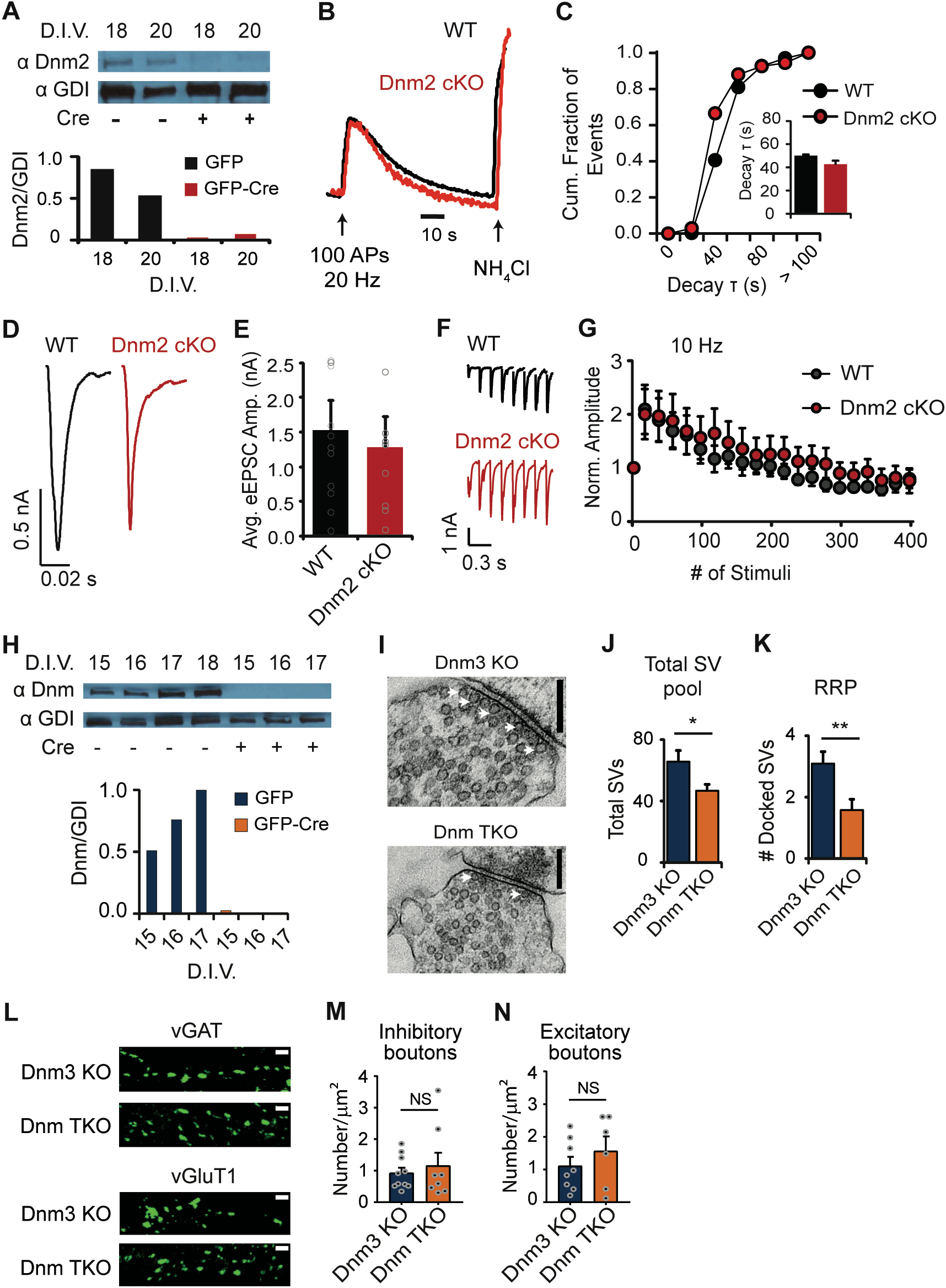
**A.** Representative Western blot (top) and quantification (bottom) of dynamin 2 levels in control (left bands) and dynamin 2 KO (right bands) neurons. There is ~93% reduction in dynamin 2 protein levels at 18-20 DIV. **B.** Example traces of normalized vGluT1-pHluorin responses to 20 Hz 100 AP (5 s) stimulation in control (WT, black) and dynamin 2 KO (red) hippocampal neuron cultures. **C.** Cumulative distribution of the calculated fluorescence decay time constant (τ). Decay time courses are fit with a single exponential function for control (WT, black) and dynamin 2 KO (red) neuronal cultures. WT: N=399 boutons; dynamin 2 KO: N=340 boutons. Inset: Average τ for WT (black) and dynamin 2 KO (red) neurons (p=0.4081 by Student’s ordinary t-test). **D.** Representative eEPSC traces from WT littermate controls (black) and dynamin 2 KO (red) cultured hippocampal neurons. **E.** Average eEPSC amplitudes from WT (N=10, mean=1.5 nA) and dynamin 2 KO (N=9, mean=1.3 nA) neurons are similar. **F.** Sample eEPSC traces of the first 8 responses from WT (black) and dynamin 2 KO (red) neurons to 400 stimuli applied at a 10 Hz frequency. **G.** Normalized responses from WT (black, N=10) and dynamin 2 KO (red, N=9) neurons after 400 pulses 10 Hz stimuli showing no effect of the absence of dynamin 2 on synaptic transmission. **H.** Representative Western blot (top) and quantification (bottom) of total dynamin (using a pan-dynamin antibody) depicting the loss of all three dynamin isoforms (Cre treated cultures, orange) compared to littermate controls (empty L307 vector, blue). There is a substantial (~93%) reduction of dynamins 1, 2 and 3 after 15 DIV in cultured hippocampal neurons. **I.** Sample electron micrograph images of Dnm 3 KO and Dnm TKO synapses. White arrows: docked synaptic vesicles. Black bars = 200 μm. **J.** Total number of synaptic vesicles (SV) per presynaptic area in Dnm 3 KO (blue, N=41 synapses, mean=65 SVs) and Dnm TKO synapses (orange, N=53 synapses, mean=47 SVs; * p= 0.0193 by Student’s ordinary t-test). **K.** Number of docked synaptic vesicles in Dnm 3 KO (blue, N=22 synapses, mean=3 SVs) and Dnm TKO synapses (orange, N=19 synapses, mean=1.5 SVs; ** p=0.0039 by Student’s ordinary t-test). These results show a significant reduction in the sizes of the total pool and the readily releasable pool of synaptic vesicles after depletion of all dynamin isoforms. **L.** Representative immunofluorescence images from Dnm 3 KO and Dnm TKO neurites stained against vGAT (top panels) and vGluT1 (bottom panels). White bars= 2 μm. **M.** Average number of inhibitory presynaptic boutons (vGAT positive) per μm^2^ in Dnm 3 KO (blue, N=10 images from 3 coverslips) and Dnm TKO cultures (orange, N=9 images from 3 coverslips). **N.** Average number of excitatory presynaptic boutons (vGluT1 positive) per μm^2^ in Dnm 3 KO (blue, N=8 images from 2 coverslips) and Dnm TKO cultures (orange, N=6 images from 2 coverslips).

We next assessed synaptic vesicle recycling in the absence of dynamin 2 after high frequency stimulation using the optical indicator of synaptic vesicle exocytosis and endocytosis vGluT1-pHluorin (Kavalali and Jorgensen, 2014; Leitz and Kavalali, 2011; Voglmaier et al., 2006). After a 20 Hz 100 action potentials (APs) stimulation, there were no significant differences in the measured decay time constant (τ) in Dnm2 cKO synapses (~46 seconds) in comparison to WT littermate control synapses (~49 seconds) (Figure 1B-C), indicating normal endocytic kinetics in neurons depleted of dynamin 2. The exocytic load was also unaltered in the absence of dynamin 2, since the ratio of the fluorescence amplitude (ΔF) after the 20 Hz stimulus to the maximal possible response obtained by perfusing ammonium chloride was similar in both groups (data not shown). We then investigated the retrieval of single synaptic vesicles after evoked exocytosis in Dnm2 cKO neurons by measuring the dwell time, the amount of time the pHluorin probe spent on the presynaptic membrane before being retrieved (Leitz and Kavalali, 2011; Chanaday and Kavalali, 2018). We observed no differences in dwell time in Dnm2 cKO neurons in comparison to the control suggesting that dynamin 2 is not required for retrieval after single synaptic vesicle release (Figure 1 - figure supplement 1A-B). In addition, amplitude of fusion events (Figure 1 - figure supplement 1C) and release probability (data not shown) were similar for control and Dnm2 cKO neurons. Finally, we assessed the effects of the loss of dynamin 2 on synaptic transmission by voltage patch clamp recordings and observed no changes in average evoked and spontaneous excitatory (eEPSC and mEPSC, respectively) activity implying that dynamin 2 is not essential for synaptic transmission (Figure 1D-G and Figure 1 - figure supplement 1D-H). Taken together, these results suggest that dynamin 2 is not critical for synaptic vesicle recycling or neurotransmission.

### Synapse morphology in the absence of dynamins 1, 2 and 3

Previous findings suggested the possibility of dynamin-independent endocytic mechanisms in neurons (Xu et al., 2008). To explore the role of all three dynamin isoforms in synaptic vesicle recycling, we crossed a knock-out (KO) mouse line for dynamin 3 and mice with floxed dynamin 1 and 2 genes. After Cre recombinase treatment, triple knock-out (TKO) neurons for all dynamin isoforms were obtained (Dnm TKO). We used hippocampal neuron cultures from littermate animals infected with the empty vector (expressing only GFP) as controls (Dnm3 KO). We used Dnm3 KOs as controls in order to obtain more consistent comparisons within littermates based on the observation that Dnm3 KOs do not show significant functional or structural deficits (Ferguson et al., 2007; Raimondi et al., 2011). To confirm that all 3 dynamin isoforms were knocked out we measured protein levels by Western blot (using a pan-dynamin antibody; Liu et al., 2008) and mRNA expression by q-RT PCR. While protein levels of all dynamins were reduced by (92.9±0.4)% (Figure 1H), mRNA levels of dynamins 1 and 2 were reduced >99% while dynamin 3 mRNA was reduced >95% in Dnm TKO neurons (Figure 1 – figure supplement 2A), indicating that all dynamins are significantly depleted under these conditions. We then analyzed synapse morphology using electron microscopy (EM) in Dnm TKO neurons and observed no gross distortions in synapse morphology at rest. However, we found a significant decrease in the total synaptic vesicle number (~65 vs 44) and the number of docked vesicles (~3 vs 1.5) in Dnm TKO neurons in comparison to Dnm3 KO littermate control neurons (Figure 1I-K). Given that neurons and synapses are only starting to develop and mature at the time of lentiviral infection, we next evaluated the effects of loss of dynamins on synapse numbers. We labeled Dnm TKO and control cultures for inhibitory synapses using an antibody against the vesicular GABA transporter (vGAT) and excitatory synapses using an antibody against the vesicular glutamate transporter (vGluT). We observed no significant differences in the total number of excitatory and inhibitory synapses per μm^2^ in Dnm TKO cultures compared to littermate controls (Figure 1L-N). This data suggests that dynamin is required for the maintenance of synaptic vesicle pool size and the docking of synaptic vesicles at rest but not for synapse formation.

### Synaptic vesicle recycling during high frequency stimulation in neurons depleted of dynamins 1,2,3

Next, we assessed synaptic vesicle endocytosis in response to 100 APs at 20 Hz (5 s) by expressing vGluT1-pHluorin in Dnm TKO neurons (Figure 2A). We observed three categorically different responses to the 20 Hz stimulus in neurons depleted of dynamins: synapses that recovered completely after 20 Hz stimulation, synapses that recovered to a baseline higher than the baseline prior to stimulation (partial recovery) and those that did not recover after stimulation (Figure 2B). To quantify this phenotype, we first calculated the amount of retrieval (ΔF_ret_) as the difference between baseline level increase after stimulation and response amplitude (ΔF), and then obtained the percentage of retrieval by dividing ΔF_ret_ by ΔF. We observed that in Dnm3 KO littermate controls, 90% of synapses recovered to 80% of the initial baseline within 150 s after stimulation, while around 60% of synapses recovered to at least 80% of the initial baseline within 150 s post stimulus in Dnm TKO neurons. This suggests that even if all three dynamin isoforms are significantly reduced, synaptic vesicles can still undergo normal endocytosis in more than half of hippocampal synapses (Figure 2B-C). We observed a decrease in the ratio of ΔF to the maximal possible response, F_NH4Cl_ (ΔF/F_NH4Cl_) in Dnm TKO compared to control (Figure 2D) suggestive of a decrease in release probability in the absence of all dynamins and corroborating our finding of decreased number of docked vesicles via EM (Figure 1K). We also detected an overall decrease in F_NH4Cl_ in Dnm TKO neurons in comparison to control suggesting a possible decrease in synaptic vesicle pool size as well (data not shown, but see Figure 1J). Finally, we estimated the kinetics of endocytosis by fitting the decay to baseline after stimulus with a single exponential decay function and used the decay time constant (τ) as a measure of endocytic rate. We observed a drastic increase in decay τ in Dnm TKO neurons indicating a slow-down in synaptic vesicle retrieval after high frequency stimulation (Figure 2E). Taken together, these results suggest that dynamins are important but not essential for synaptic vesicle recycling during high frequency neuronal activity.

**Figure 2.**
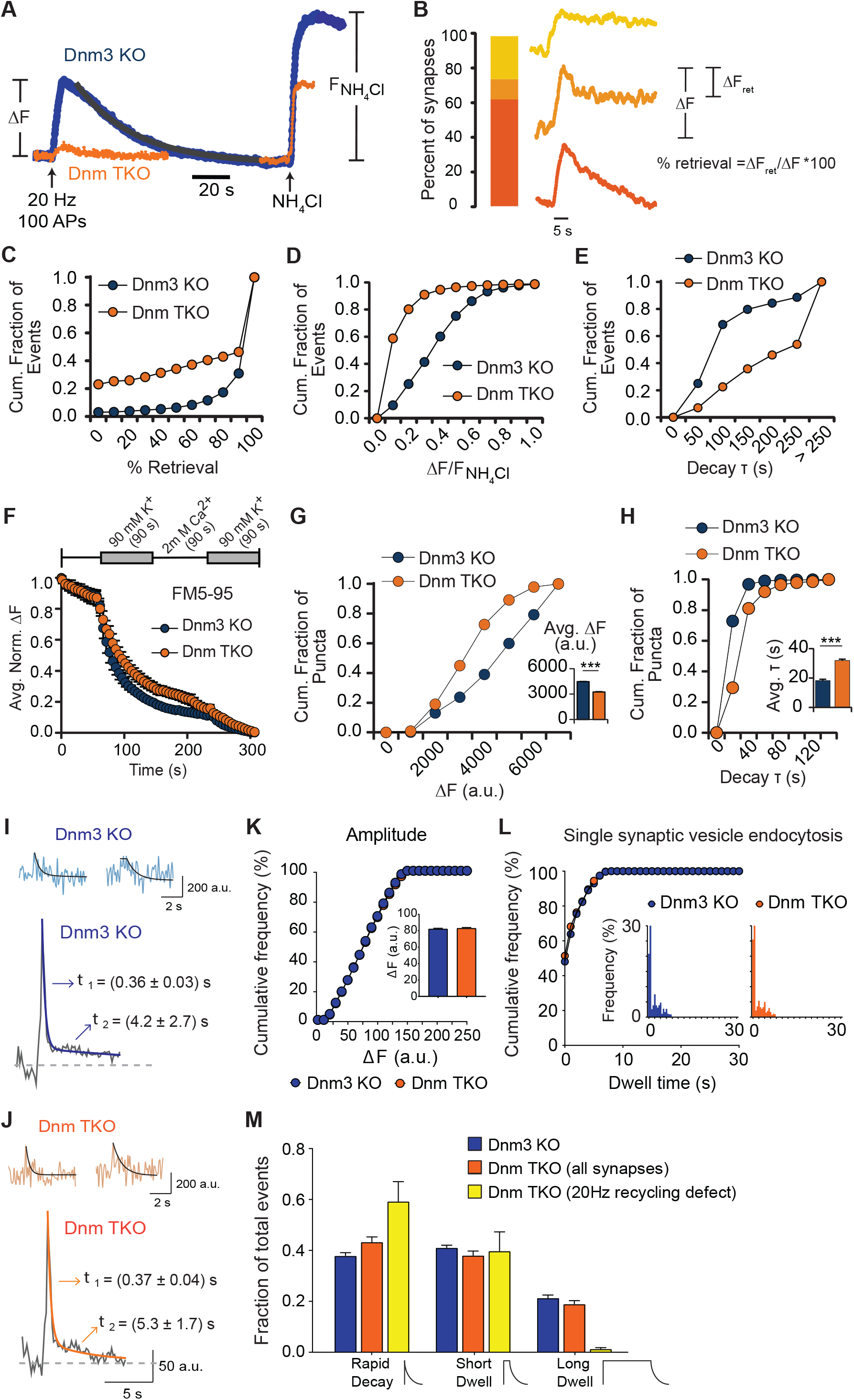
**A.** Sample vGluT1-pHluorin traces in Dnm 3 KO (blue) and Dnm TKO (orange) boutons in response to 20 Hz 100 AP (5 s) stimulation (ΔF) and after perfusion of 20 mM NH_4_Cl (F_NH4Cl_). **B.** Percent distribution of the different retrieval types observed in Dnm TKO neurons following 20 Hz stimulation. 63% of Dnm TKO boutons decayed back to at least 70% of the initial increase in fluorescence (ΔF) indicating high (similar to control) retrieval of the fused synaptic vesicle proteins. 25% of Dnm TKO synapses had obstructed endocytosis since they decayed to less than 20% of the initial increase in fluorescence while the remaining 12% of Dnm TKO synapses showed partial variable efficiency in retrieval with decays between 20% - 70% of the initial increase in fluorescence. **C.** Cumulative histogram of vGluT1-pHluorin retrieval (% of ΔF_ret_/ΔF) after 20 Hz 100 AP stimulation for Dnm 3 KO (blue, N=1885 boutons) and Dnm TKO (orange, N=967 boutons) showing an impairment in retrieval efficiency in the absence of all dynamin isoforms (p<0.0001, Kolmogorov-Smirnov test). **D.** Cumulative distribution of response amplitudes (ΔF relative to F_NH4Cl_) after 20 Hz 100 AP stimulation for Dnm 3 KO (blue, N=1885 boutons) and Dnm TKO (orange, N=967 boutons) revealing a decrease of exocytosis in the absence of all dynamin isoforms (p<0.0001, Kolmogorov-Smirnov test). **E.** Cumulative distribution of calculated decay time constant (τ) for Dnm 3 KO (blue, mean=85 s) and Dnm TKO (orange, mean=137 s) showing a slowdown in endocytosis in neurons lacking dynamins 1, 2 and 3 (p<0.0001, Kolmogorov-Smirnov test). **F.** Top: outline of FM dye release experiment. Synapses were preloaded with FM dye by incubating cultures in 45 mM KCl modified Tyrode’s buffer containing 18 μM FM5-95. Release of dye was induced by incubating cultures in 90 mM KCl modified Tyrode’s buffer. Bottom: Average normalized ΔF response in Dnm 3 KO (blue, N=5 coverslips) and Dnm TKO (orange, N=5 coverslips) synapses after two successive stimulations with 90 mM KCl to induce release of FM5-95. **G.** Cumulative distribution of the total change in fluorescence (ΔF) after 2 cycles of 90 mM KCl stimulation and FM5-95 dye release in Dnm 3 KO (blue) and Dnm TKO (orange) synapses showing a significant decrease in exocytosis in the absence of dynamins (*** p<0.0001, Kolmogorov-Smirnov test). Inset: average values of ΔF after 2 cycles of 90 mM KCl stimulation. **H.** Cumulative histogram of the decay time constant (τ) from fitting FM5-95 fluorescence decay after the first 90 mM KCl stimulation, in Dnm 3 KO (blue) and Dnm TKO (orange) presynaptic boutons showing a slowdown in the kinetics of exocytosis in neurons lacking all dynamin isoforms (*** p<0.0001 by Kolmogorov-Smirnov test). Inset: average τ values. **I.** Analysis of single vesicle fusion events in Dnm 3 KO synapses. Top: two representative single vesicle fusion events monitored with vGluT1-pHluorin (black lines show the dwell time and the subsequent exponential decay of fluorescence). Bottom: average of all single vesicle event traces fitted with a double exponential decay (black line) revealing the two components of endocytosis (ultrafast ~360 ms; fast ~8.1 s). **J.** Analysis of single vesicle fusion events in Dnm TKO synapses. Top: two representative single vesicle fusion events monitored with vGluT1-pHluorin (black lines show the dwell time and the subsequent exponential decay of fluorescence). Bottom: average of all single vesicle event traces fitted with a double exponential decay (black line) revealing the two components of endocytosis (ultrafast ~360 ms; fast ~6.7 s). There is not a major impact of removal of all dynamin isoforms in the kinetics of single synaptic vesicle retrieval and reacidification during low frequency stimulation. **K.** Cumulative histogram of amplitudes for single synaptic vesicle fusion events in Dnm 3 KO (blue, N=1565 boutons) and Dnm TKO (orange, N=961 boutons) show a similar distribution between both groups. Inset: average amplitude (ΔF) for Dnm 3 KO (blue, mean=82.2 a.u.) and Dnm TKO (orange, mean=83.0 a.u., p=0.9137 by Kolmogorov-Smirnov test). **L.** Cumulative distribution of dwell times for Dnm 3 KO (blue, N=1565 boutons) and Dnm TKO (orange, N=961 boutons). The lack of all three dynamin isoforms does not impact the kinetics of single synaptic vesicle endocytosis. Inset: frequency histogram of dwell time duration for each experimental group (p=0.06752 by Kolmogorov-Smirnov test). **M.** Classification of dwell times into three categories: Rapid decay (<1 s dwell), Short decay (1 – 10 s dwell) and Long dwell (> 10 s dwell). The fraction of total measured dwell times into each category are shown for Dnm 3 KO (blue) and Dnm TKO (orange), showing no differences in the proportion of fast or slow endocytic events between the groups. When only events from synapses with defects in endocytosis after high frequency stimulation (yellow traces in B, yellow bars in M) were considered for the analysis, no significant defects in dwell times (single synaptic vesicle endocytosis) were observed (yellow).

Given our observation of almost normal synaptic vesicle endocytosis for ~60% of synapses after 20 Hz stimulation, we next asked whether the retrieved vesicles are available for re-release. We stimulated synaptic uptake of FM5-95 dye by incubating Dnm TKO and littermate control neurons in 45 mM K^+^ modified Tyrode’s solution containing 18 μM FM5-95 dye for 2 minutes. Excess dye was washed out, and then we applied a second stimulation to trigger synaptic vesicle exocytosis and dye release by incubating neuronal cultures in 90 mM K^+^ modified Tyrode’s solution for 90 s, two consecutive times (Figure 2F). We observed a decrease in fluorescence indicative of dye release in both littermate control and Dnm TKO synapses (Figure 3F). However, the decrease in fluorescence in response to 90 mM K^+^ stimulation was greater in littermate controls than in Dnm TKO (Figure 2G) and it also occurred with a slower time-course in Dnm TKO synapses (Figure 2H). These results suggest that depletion of dynamins reduces the total pool size of synaptic vesicles and the remaining vesicles show slower mobilization and recovery compared to controls during strong depolarization. Our findings are also consistent with a function for dynamins in the regulation of synaptic vesicle release probability (Lou et al., 2012 and see below).

**Figure 3.**
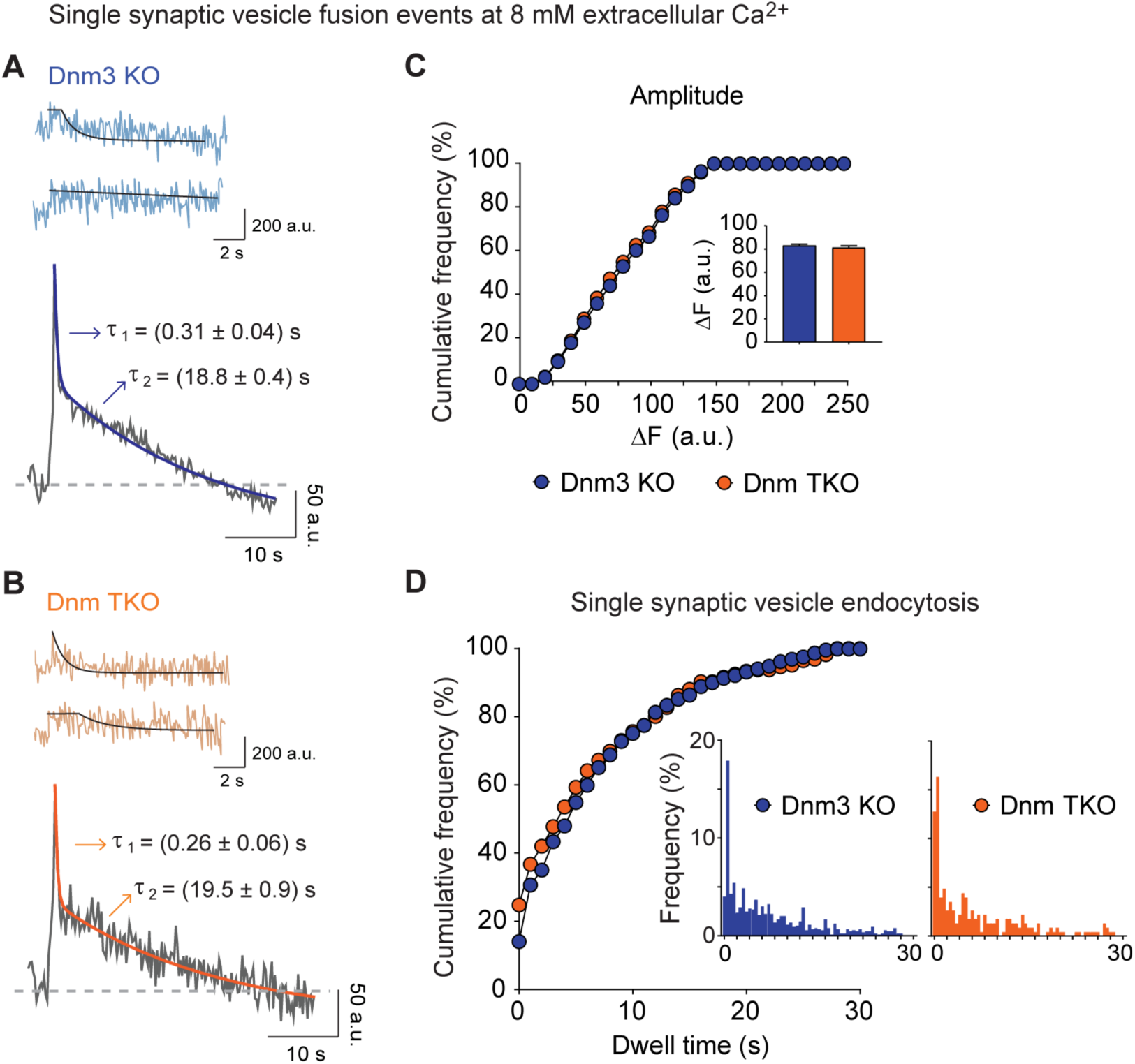
**A.** Analysis of single vesicle fusion events in Dmn 3 KO synapses at extracellular 8 mM Ca^2+^ concentration. Top: two representative single vesicle fusion events (blue) monitored with vGluT1-pHluorin (black lines show the dwell time and the subsequent exponential decay of fluorescence). Bottom: average of all single vesicle event traces fitted with a double exponential decay (blue line) revealing the two components of endocytosis (ultrafast ~310 ms; fast ~19 s; N=888 boutons). **B.** Analysis of single vesicle fusion events in Dmn TKO synapses at extracellular 8 mM Ca^2+^ concentration. Top: two representative single vesicle fusion events (orange) monitored with vGluT1-pHluorin (black lines show the dwell time and the subsequent exponential decay of fluorescence). Bottom: average of all single vesicle event traces fitted with a double exponential decay (orange line) revealing the two components of endocytosis (ultrafast ~260 ms; fast ~19 s; N=226 boutons). There is not a major impact of removal of all dynamin isoforms in the kinetics of single synaptic vesicle retrieval and reacidification during low frequency stimulation. Moreover, in the absence of dynamins the Ca^2+^-dependent slowdown of endocytosis occurs normally. **C.** Cumulative distribution of single vesicle event amplitudes from Dnm 3 KO (blue, N=888 boutons) and Dnm TKO (orange, N=226 boutons) synapses in 8mM Ca^2+^. Inset: Average amplitude for Dnm 3 KO (blue, mean=83.0 a.u.) and Dnm TKO (orange, mean=81.2 a.u., p=0.8619 by Kolmogorov-Smirnov test) are similar between the two experimental groups. **D.** Cumulative distribution of single vesicle event dwell times from Dnm 3 KO (blue, N=888 boutons) and Dnm TKO (orange, N=226 boutons) synapses in extracellular 8mM Ca^2+^. There is no effect in the kinetics of single synaptic vesicle endocytosis after removal of all dynamin isoforms. Inset: Histograms of dwell time duration for both experimental groups (p=0.1149 by Kolmogorov-Smirnov test).

### Single synaptic vesicle endocytosis is dynamin independent

So far, we have described dynamin function in synaptic vesicle recycling during strong stimulation. To address the importance of dynamins in the recycling of individual synaptic vesicles, we applied sparse, single AP stimulation to Dnm TKO hippocampal neuron cultures expressing vGluT1-pHluorin. We observed no change in the amplitude of single synaptic vesicle fusion events in Dnm TKO synapses in comparison to littermate control Dnm3 KO synapses (Figure 2I-K). No differences were observed in the distribution of dwell times in presynaptic boutons from both groups (Figure 2L). Moreover, exponential fitting of the ensemble average traces from Dnm3 KO and TKO groups revealed the presence of two phases of decay, one ultrafast (~300 ms) and one fast (~6-8 s) of similar characteristics for both groups (Figure 2I-J) (see Chanaday and Kavalali, 2018). This further supports the absence of defects in retrieval and re-acidification in dynamin TKO neurons compared to Dnm3 KO. Finally, we categorized the single synaptic vesicle fusion events into three groups based on the event profile (see sample traces in Figure 2I-J). Rapid decay were events that decayed almost instantaneously after fusion (<500 ms) and may correspond to ultrafast endocytosis events. Short dwell were events that resided on the membrane for more than 500 ms and decayed to baseline before the end of the allotted timeframe (500 ms – 10 s). Long dwell were events that did not decay to baseline within the allotted time window (> 10 s). We measured the fractional composition of the single synaptic vesicle fusion events in these three categories and noted no changes in the distribution between Dnm TKO and littermate control Dnm 3 KO synapses (Figure 2M). Long dwell events, which are indicative of protein stranded at the plasma membrane and not retrieved during the imaging window, constituted 19±1% of total dynamin TKO single vesicle events in comparison to 21±1% of total control single vesicle events (Figure 2M). We also looked at the fractional composition of single synaptic vesicle events from Dnm TKO synapses with severe retardation of synaptic vesicle endocytosis after high frequency stimulation and observed that only ~1% of single vesicle events from these synapses were long-dwell events. This observation suggests that the negative impact of dynamin loss-of-function in these synapses was limited to multi-vesicle endocytosis and the dynamin-regulation of single vesicle versus multi-vesicle retrieval events are relatively independent (Figure 2M). We assessed the distribution of rapid decay and short dwell events and observed no differences between Dnm 3 KO and Dnm TKO (Figure 2M). Taken together, our data suggest that the kinetic properties of single synaptic vesicle retrieval are largely dynamin independent.

### Dynamins are not essential for Ca^2+^ dependent delay in single synaptic vesicle retrieval

Prior work in our lab has demonstrated that increasing extracellular Ca^2+^ can slow down endocytosis after single synaptic vesicle release (Leitz and Kavalali, 2011). This retardation in synaptic vesicle retrieval is dependent on synaptotagmin-1 (syt1), the Ca^2+^ sensor for synchronous synaptic vesicle exocytosis (Li et al., 2017; Chanaday and Kavalali, 2018). In chromaffin cells, syt1 has been shown to interact directly with the PH domain of dynamin 1 (Dnm1) to regulate the fission pore of single dense core vesicles (McAdam et al., 2015). To address the role of dynamins in the modulation of synaptic vesicle retrieval kinetics by Ca^2+^, we increased extracellular Ca^2+^ concentration to 8 mM and evoked single synaptic vesicle release in Dnm TKO and littermate control neurons (Figure 3A-B). We observed no differences in the amplitude of single synaptic vesicle fusion events in these conditions (Figure 3C). The distribution of dwell times (Figure 3D) as well as the two components of fluorescence decay for the average traces (Figure 3A-B) were similar for Dnm3 KO and Dnm TKO synapses. This result indicates that Ca^2+^ can slow down single synaptic vesicle endocytosis in synapses lacking all dynamin isoforms, as described before for wild-type neurons (Leitz and Kavalali, 2011; Li et al., 2017) leading us to conclude that dynamins are not essential for Ca^2+^-dependent prolongation of single synaptic vesicle dwell time. Moreover, based on the analysis of the ensemble averaged fluorescent traces, the kinetics of ultrafast endocytosis and fast endocytosis components of retrieval do not seem to be modulated by dynamins (Chanaday and Kavalali, 2018).

### Inhibitory neurotransmission after depletion of dynamins

Our data suggests that the postnatal deletion of dynamin results in a decreased pool size of synaptic vesicles that can recycle with sparse stimulation but are limited upon high frequency stimulation. Prior work has demonstrated that the efficiency of synaptic vesicle endocytosis is dependent on the preceding exocytic load (Armbruster et al., 2013). Inhibitory synapses have been reported to undergo tonic neurotransmission and as such require more efficient endocytic mechanisms to maintain fidelity of neurotransmission (Evergren et al., 2006; Swadlow, 2003). Therefore, synaptic vesicle recycling in non-glutamatergic synapses might be different from glutamatergic synapses. To assess the dynamin dependence of inhibitory transmission, we recorded evoked inhibitory postsynaptic currents (eIPSC) in Dnm TKO neurons (Figure 4). We observed a decrease in the average eIPSC amplitude (0.8 nA vs 1.3 nA) in Dnm TKO neurons in comparison to the littermate controls (Figure 4A). We applied a train of 30 pulses at 1 Hz and observed similar levels of synaptic depression in Dnm TKO and littermate control cultures (Figure 4B-C). Similarly, we observed similar levels of synaptic depression after we challenged Dnm TKO and littermate control neurons with a 400 pulse 10 Hz train stimulus (Figure 4D-E). However, when we stimulated with 1000 pulses at 30 Hz frequency, we observed an initial synaptic facilitation in Dnm TKO neurons that was absent in littermate control Dnm3 KO neurons. This data is consistent with the premise that the absence of dynamin in inhibitory neurons leads to a decrease in release probability. We also investigated spontaneous or miniature inhibitory neurotransmission (mIPSC) in the presence of tetrodotoxin (TTX). We observed no significant changes in the frequency of mIPSC events between groups (1.8 Hz vs 1.4 Hz) but found a rightward shift in the cumulative distribution of amplitudes for Dnm TKO neurons indicative of an overall increase in mIPSC amplitude (Figure 4G-I). Since no alterations in synaptic vesicle size were found by EM (data not shown), this could indicate differences in postsynaptic levels of GABA receptors. In conclusion, for inhibitory synapses, depletion of all three dynamin isoforms leads to a reduction of evoked release probability and an increase in amplitude of spontaneous postsynaptic responses.

**Figure 4.**
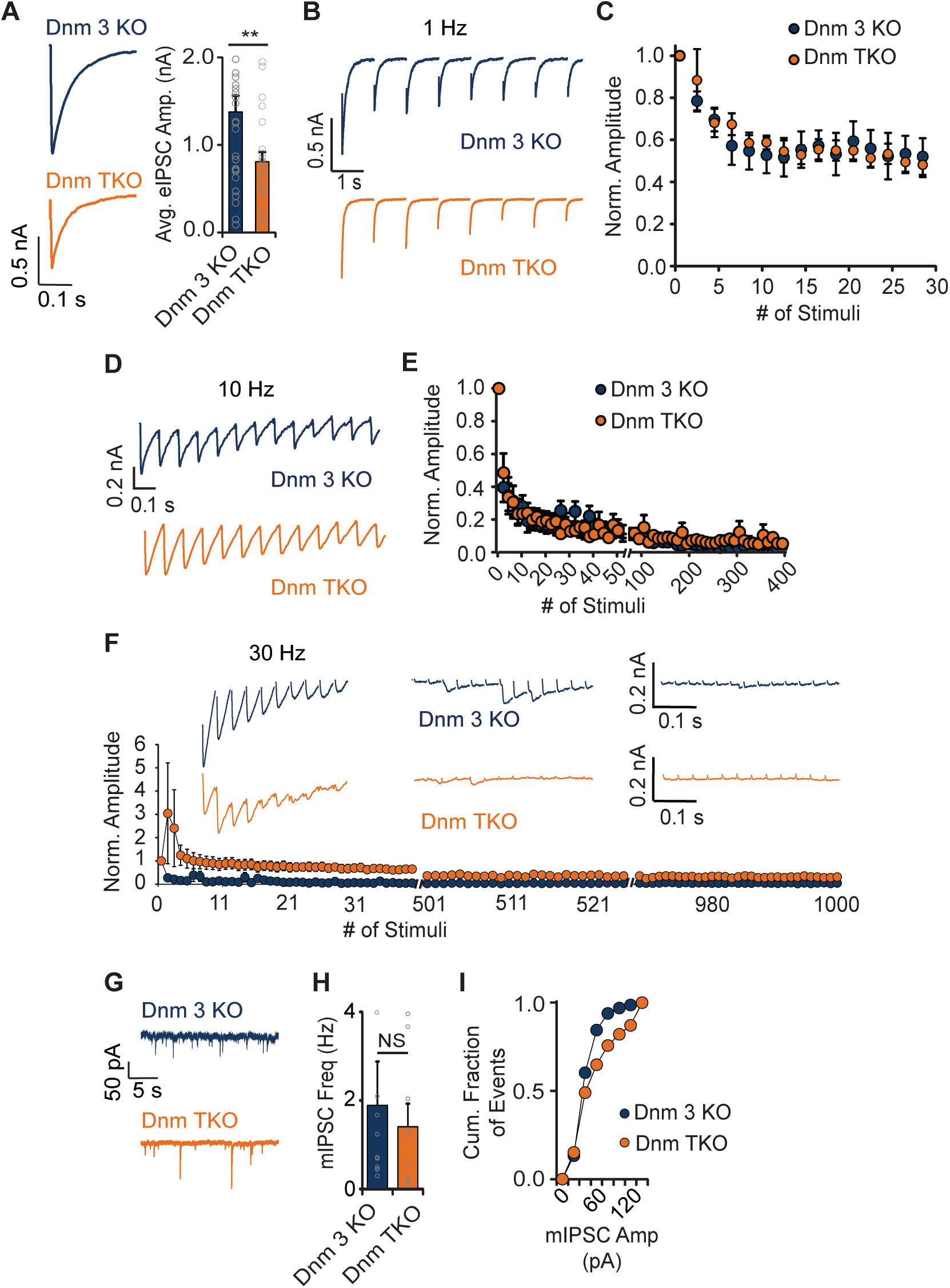
**A.** Left, sample eIPSC traces after a single stimulus from littermate control Dnm 3 KO (blue) and Dnm TKO (orange) neurons. Right, average eIPSC amplitudes of littermate control Dnm 3 KO (blue, mean=1.4 nA, N=31) and Dnm TKO (orange, mean =0.8 nA, N=27) hippocampal neuron cultures (** p = 0.0172, Student’s ordinary t-test). There is a decrease in amplitude in Dnm TKO neurons indicating a reduction in release probability. **B.** Representative traces of the first 8 eIPSC responses to a 1 Hz 30 AP stimulus for Dnm 3 KO (blue) and Dnm TKO (orange). **C.** Normalized eIPSC amplitudes after 1 Hz 30 AP stimulus for Dnm 3 KO (blue, N=9) and Dnm TKO (orange, N=12) showing no differences between the groups (p=0.9917, Two-way RM ANOVA). **D.** Sample traces of the first 13 eIPSC responses to a 10 Hz 400 AP stimulus for Dnm 3 KO (blue) and Dnm TKO (orange) neurons. **E.** Normalized eIPSC amplitudes after 10 Hz 400 AP stimulus for Dnm 3 KO (blue, N=8) and Dnm TKO (orange, N=12) revealing similar depression time course for both groups (p=0.7167, Two-way RM ANOVA). **F.** Top: sample traces of the first 10, middle 12 and last 12 eIPSC responses to a 30 Hz 1000 AP stimulus for Dnm 3 KO (blue) and Dnm TKO (orange). Bottom: normalized eIPSC amplitudes after 30 Hz 1000 AP stimulus for Dnm 3 KO (blue, N=11) and Dnm TKO (orange, N=7). There is a facilitation of release at the beginning of the stimulation train in Dnm TKO indicative of reduced release probability in this group (p=0.0017, Two-way RM ANOVA). **G.** Representative mIPSC recordings from Dnm 3 KO (blue) and Dnm TKO (orange) neurons. **H.** Average mIPSC frequency for Dnm 3 KO (blue, N=10, mean=1.8 Hz) and Dnm TKO (orange, N=11, mean=1.4 Hz) hippocampal neuron cultures reveals no effect on the rate of inhibitory spontaneous neurotransmitter release after depletion of all dynamins (p=0.6636, Student’s ordinary t-test). **I.** Cumulative distribution of the mIPSC amplitudes for Dnm 3 KO (orange) and Dnm TKO (blue) showing a small but significant reduction in amplitudes (p<0.0001, Kolmogorov Smirnov test).

### Loss of dynamins 1, 2 and 3 impairs excitatory neurotransmission

Excitatory neurotransmission was previously shown to have reduced release probability and a decrease in the amplitude of postsynaptic currents in the absence of the neuronal dynamin isoforms, dynamin 1 and 3. These alterations could be reversed by chronic suppression of activity (Lou et al., 2012) indicating that it results from vesicle depletion due to the endocytic arrest. To evaluate excitatory neurotransmission when all dynamin isoforms are depleted, we measured evoked excitatory postsynaptic currents (eEPSC) in Dnm TKO hippocampal neurons (Figure 5). We observed a significant decrease in eEPSC amplitude (1.1 nA vs 0.1 nA) in Dnm TKO neurons (Figure 5A) accompanied by facilitation to a train of 1 Hz and 10 Hz stimuli (Figure 5B-E) suggestive of a decrease in the probability of release in Dnm TKO neurons compared to Dnm3 KO littermate control. We also investigated changes in spontaneous neurotransmission in the presence of TTX in Dnm TKO neurons. We observed a decrease in mEPSC frequency (1.7 Hz vs 0.7 Hz) and an increase in mEPSC amplitude (Figure 5F-H) in Dnm TKO neurons compared to control. These findings are consistent with prior reports assessing excitatory neurotransmission defects in constitutive dynamin 1,3 double KO neurons (Lou et al., 2012) pointing to a negligible role for dynamin 2 in excitatory neurotransmission.

**Figure 5.**
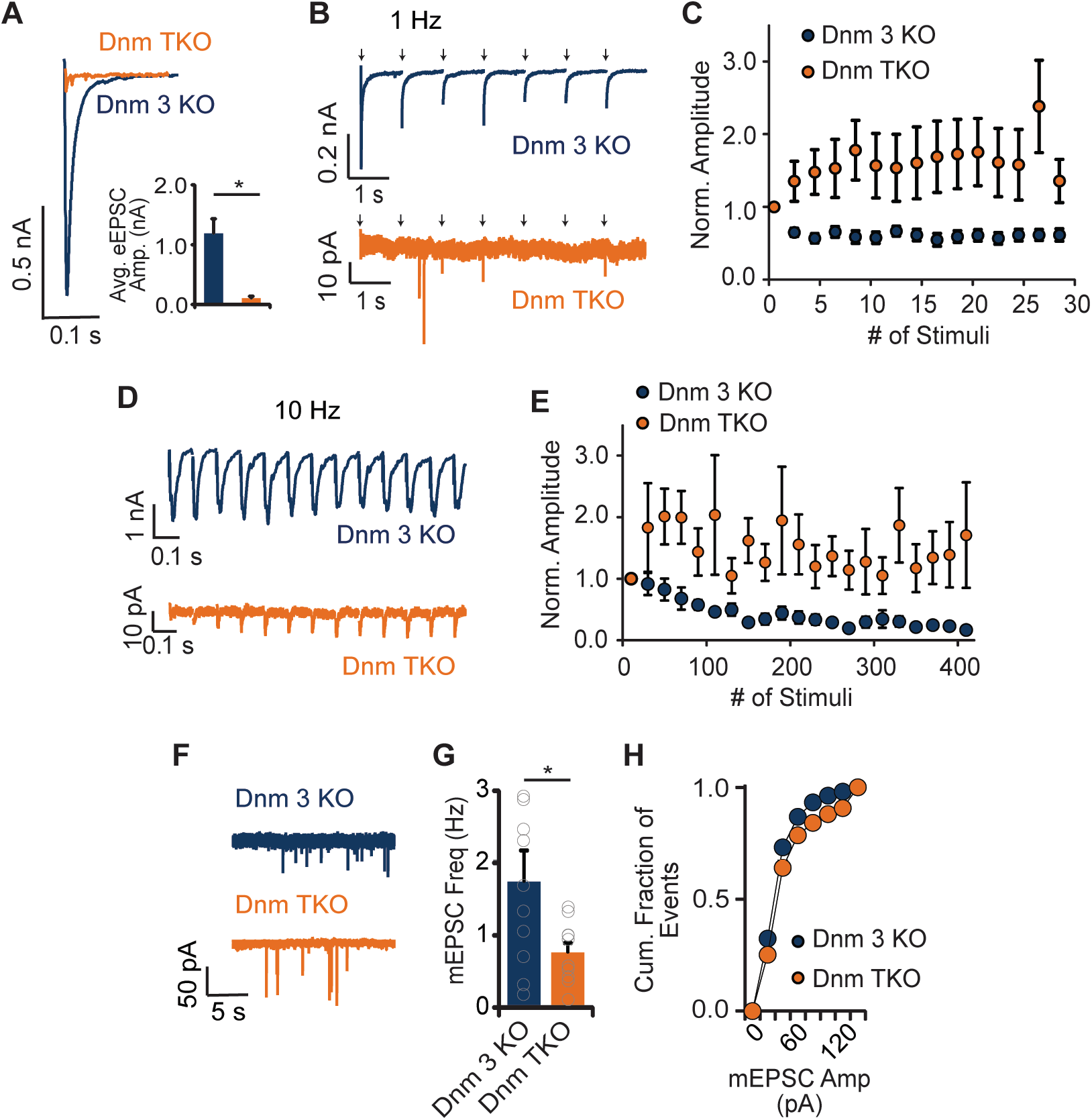
**A.** Sample eEPSC traces after a single stimulus from littermate control Dnm 3 KO (blue) and Dnm TKO (orange) neurons. Inset: Average eEPSC amplitudes of littermate control Dnm 3 KO (blue, mean=1.2 nA, N=10) and Dnm TKO (orange, mean=0.1 nA, N=10) neuronal cultures revealing a severe reduction in release probability at excitatory synapses (* p=0.0003, Student’s ordinary t-test). **B.** Representative traces of the first 7 eEPSC responses to a 1 Hz 30 AP stimulus for Dnm 3 KO (blue) and Dnm TKO (orange). Arrows indicate application of stimulus, note the abnormal presence of failures in Dnm TKO trace. **C.** Normalized eEPSC amplitudes after 1 Hz 30 AP stimulus for Dnm 3 KO (blue, N=10) and Dnm TKO (orange, N=10) showing facilitation in Dnm TKO consistent with reduced release probability (p = 0.0064, Two-way RM ANOVA). **D.** Sample traces of the first 13 eEPSC responses to a 10 Hz 400 AP stimulus for Dnm 3 KO (blue) and Dnm TKO (orange). **E.** Normalized eEPSC amplitudes after 10 Hz 400 AP stimulus for Dnm 3 KO (blue, N=7) and Dnm TKO (orange, N=8), depletion of all dynamin isoforms has a severe negative effect in the probability of release at excitatory synapses (p = 0.0182, Two-way RM ANOVA). **F.** Representative mEPSC recordings from Dnm 3 KO (blue) and Dnm TKO (orange) neurons. **G.** Average mEPSC frequency for Dnm 3 KO (blue, N=10, mean=1.74 Hz) and Dnm TKO (orange, N=11, mean=0.76 Hz) hippocampal neuron cultures revealing a >2-fold reduction in the rate of spontaneous release (* p=0.0353, Student’s ordinary t-test). **H.** Cumulative distribution of mEPSC amplitudes for Dnm 3 KO (blue) and Dnm TKO (orange) showing no significant difference between the groups (p<0.0001, Kolmogorov-Smirnov test).

### Spontaneously fused synaptic vesicles recycle independently of actin, DRP-1 and Arp2/3 complex in dynamin TKO neurons

Our results so far have revealed that dynamin is not required for the recycling of a single synaptic vesicle after either evoked or spontaneous fusion. We next attempted to identify crucial proteins for the recycling of a single synaptic vesicle in Dnm TKO neurons. In multiple cell types, dynamin is localized to Arp2/3 complex enucleated actin meshwork (Baldassarre et al., 2003; Gold et al., 1999; Lee and De Camilli, 2002; Orth et al., 2002; Schlunck et al., 2004). Similarly, during clathrin-mediated endocytosis, both actin and dynamin are recruited to the site of retrieval in the early phase suggesting a functional significance in initiation and maturation of clathrin-coated pits (Grassart et al., 2014). In both the large calyx of Held and small central hippocampal synapses, knockout of actin isoforms revealed that actin functions in all forms of synaptic vesicle endocytosis (Wu et al., 2016). Here, we explored the role of actin and Arp2/3 complex in the recycling of single synaptic vesicles by using the small molecule inhibitors Latrunculin A and CK-666 to inhibit actin and Arp2/3 complex, respectively. We also assessed dynamin-related protein 1 (DRP-1) function in synaptic vesicle recycling using the specific inhibitor Mdivi-1, as previous studies described that mutations in DRP in *Drosophila* leads to synaptic vesicle depletion (Rikhy et al., 2007). We knocked out all isoforms of dynamin in cultured hippocampal neurons and recorded mEPSC events before and after acute (10 min) treatment with each small molecule inhibitor (Figure 6). These experiments did not reveal significant differences in mEPSC frequency after treatment with Latrunculin A, CK-666 or Mdivi-1 (Figure 6A-B). Taken together, our findings suggest that dynamin, actin, arp2/3 complex and DRP-1 do not play a major role in the recycling of single spontaneously released synaptic vesicles.

**Figure 6.**
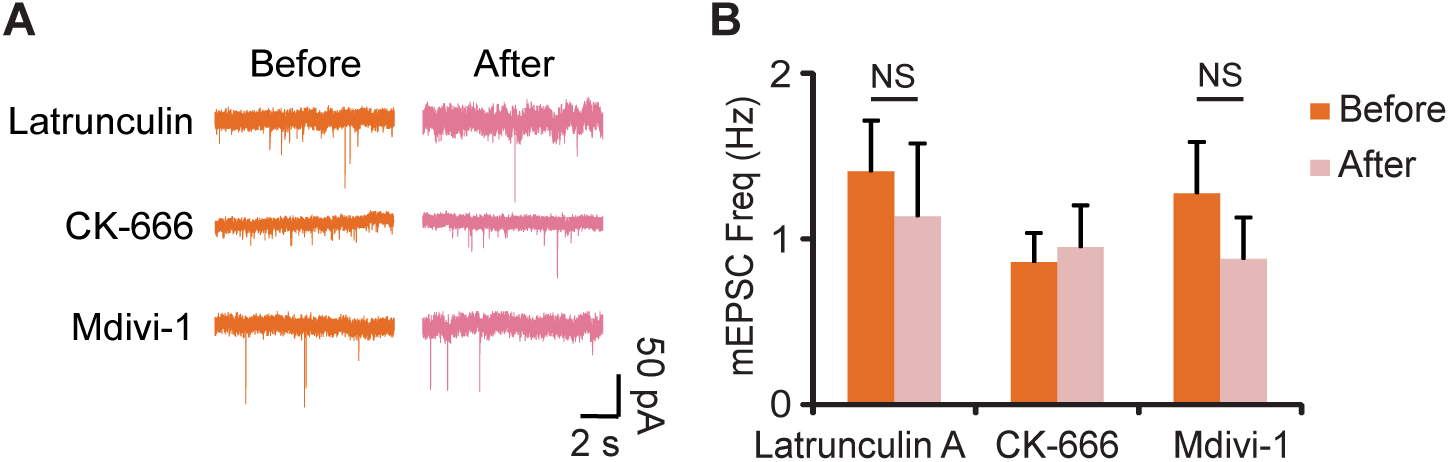
**A.** Sample mEPSC traces from Dnm TKO hippocampal neuron cultures recorded before (orange) and after (pink) treatment with Latrunculin A (20 μM), CK-666 (200 μM) and Mdivi-1 (50 μM). **B.** Average mEPSC frequency from Dnm TKO neurons before (orange) and after (pink) treatment with Latrunculin A (N=9, p=0.4856, Student’s paired t-test), CK-666 (N=6, p=0.7339, Student’s paired t-test) and Mdivi-1 (N=8, p=0.3597, Student’s paired t-test). Actin dynamics, Arp2/3 and the mitochondrial dynamin-related protein (DRP) do not mediate the remaining excitatory neurotransmission after depletion of all dynamin isoforms.

## Discussion

In this study, to investigate the functional diversity of dynamins, we established a colony of dynamin mutant mice comprised of dynamin-1 and dynamin-2 conditional knockouts (expressing floxed alleles of dynamin-1 and -2), and dynamin-3 constitutive knockouts. Using conditional knockouts to suppress dynamin expression in vitro, we could avoid systemic effects and impair dynamin function after synaptogenesis thus circumventing major developmental effects. When we assessed dynamin-2 loss-of-function in synaptic vesicle recycling and neurotransmission, we did not observe any apparent defects after the postnatal deletion of dynamin-2. In contrast, depletion of all three dynamins (dynamin-1, -2 and -3) in hippocampal neurons in culture impaired neurotransmission, including a decrease in release probability and also a substantially slowed endocytosis after high frequency stimulation (20 Hz). These effects were more pronounced in recordings of glutamatergic synaptic transmission compared to GABAergic neurotransmission suggesting a potential difference in dynamin-dependence between the two neurotransmitter systems. We also visualized dynamin TKO synapses via electron microscopy and observed decreases in both synaptic vesicle number and number of docked vesicles, which may in part account for the decrease in evoked neurotransmitter release probability.

Despite the apparent decrease in evoked release, the frequency of spontaneous neurotransmitter release events (as detected by postsynaptic voltage-clamp recordings) and the kinetics of single vesicle fusion-retrieval events (as detected optically) were only mildly affected. This surprising result is, nevertheless, consistent with earlier findings based on alternative methods from our group as well as others. For instance, treating neurons with the small molecule inhibitor of dynamin, dynasore, capable of inhibiting endocytosis, did not elicit any defects in spontaneous neurotransmission although evoked synchronous and asynchronous transmission showed significant activity-dependent suppression (Chung et al., 2010). Similarly, the injection of non-hydrolyzable GTP into the calyx of Held synapse revealed an initial block of synaptic vesicle retrieval that was followed by a resumption of synaptic vesicle endocytosis (Xu et al., 2008). In addition, a study in salamander retinal cone cells, a tonically active synapse, revealed no change in endocytosis in the presence of dynamin inhibitors (Van Hook and Thoreson, 2012). These reports suggest the presence of dynamin-independent endocytosis mechanisms in neurons contributing to the retrieval of synaptic vesicles from the presynaptic plasma membrane.

Our results suggest that dynamins play a key role in regulation of evoked probability of release and the retrieval of synaptic vesicle components after strong stimulation. In addition to the decrease in synaptic vesicle numbers, decreased release probability can be a consequence of limited availability of release sites due to imperfect clearance of fused vesicles during sustained high frequency activity (Kawasaki et al., 2000; Lou et al., 2012; Hua et al., 2013). Given the relative sparsity of spontaneous fusion events, the impairments in clearance of release sites may not present a major impediment for spontaneous fusion propensity. However, it is also important to note that in the absence of dynamins we did not detect a major impairment in the kinetic properties of retrieval after single synaptic vesicle fusion events indicating that the maintenance of spontaneous fusion events cannot solely be explained by their low frequency, but rather suggest a role for dynamin-independent recycling mechanisms.

In this study, we used lentiviral delivery of Cre-recombinase to delete dynamin-1 and dynamin-2 expression on the background of dynamin-3 constitutive knock out neurons. This approach resulted in swift depletion of dynamins as detected by Western blots as well as quantitative RT-PCR analysis. This substantial depletion of all dynamin isoforms is consistent with earlier measurements of dynamin protein lifetime in neurons (Fornasiero et al., 2018). However, we cannot fully exclude the possibility that some residual dynamins, below our detection, may contribute to the intact synaptic vesicle trafficking events we detect under these conditions. Such an effect of residual dynamins would nevertheless suggest a steep functional selectivity of dynamins depending on their abundance. According to this premise, while ultralow dynamin levels would be sufficient to maintain quantal synaptic vesicle trafficking, majority of activity driven synaptic vesicle endocytosis requires high levels of dynamin expression.

These results open new critical questions. What are the possible mechanisms that activate this putative dynamin-independent endocytosis that appears to be specific to single synaptic vesicle retrieval and spontaneous neurotransmission? How does membrane scission operate without the perennial pinchase dynamin? Although we do not have answers to these questions at this time, our findings highlight some possibilities. Our findings using small molecule inhibitors suggest that actin is not a key player for maintenance of quantal neurotransmission when dynamins are depleted. Recent studies focusing on the uptake of bacterial Shiga and cholera toxins, revealed that clathrin-independent endocytic events — that are often less reliant on dynamin — actually utilize BAR domain proteins such as endophilins for membrane scission (Renard et al., 2015). Therefore, we cannot exclude the possibility that endophilins such as endophilin-A2 and the synapse enriched endophilin-A1 (e.g. Llobet et al., 2011) may play critical roles in dynamin-independent synaptic vesicle retrieval. Overall, future experiments aimed at uncovering the molecular mechanism of single synaptic vesicle retrieval will also help elucidate the physiological role of this dynamin-independent quantal neurotransmission.

## Materials and Methods

### KEY RESOURCES TABLE

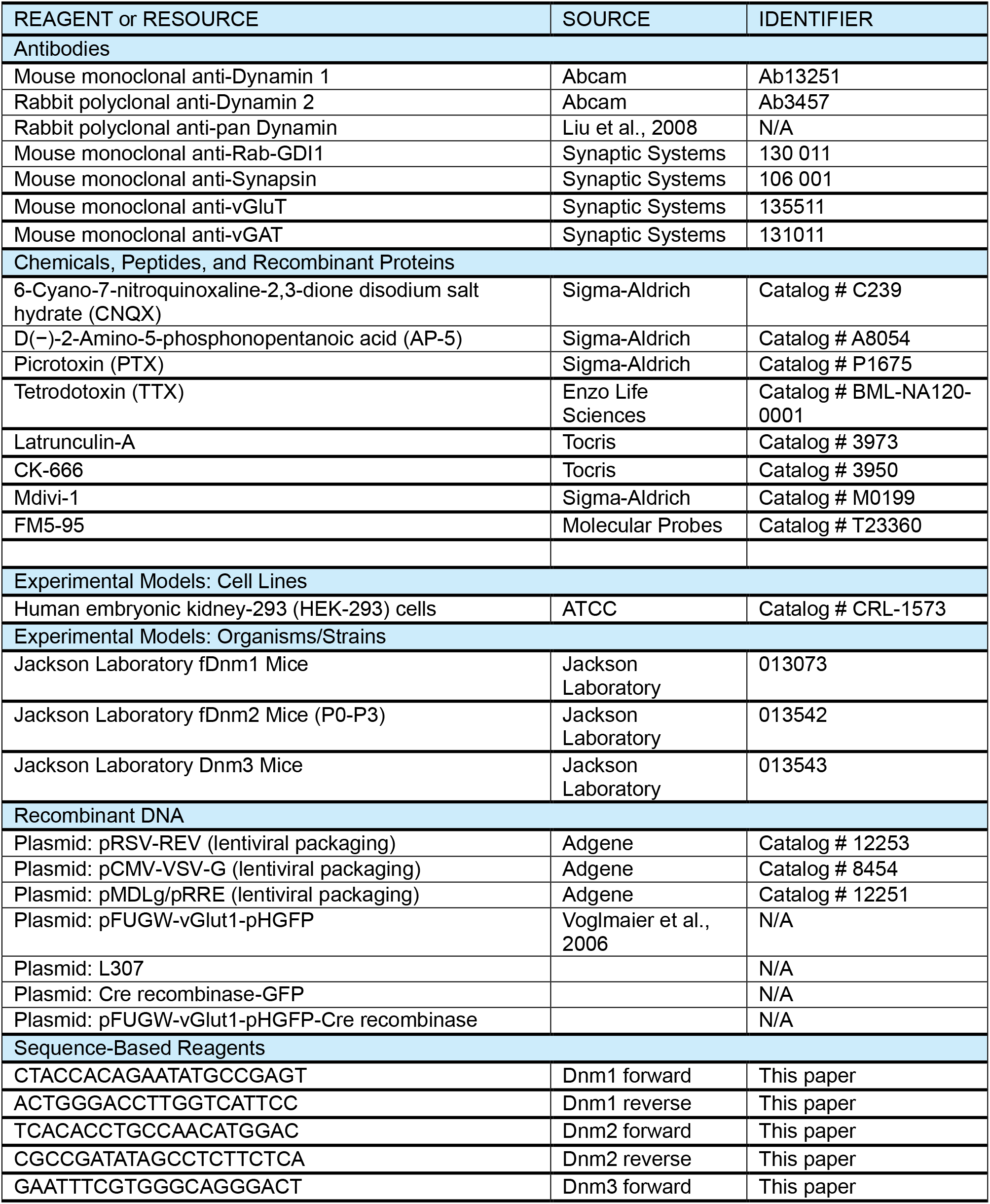
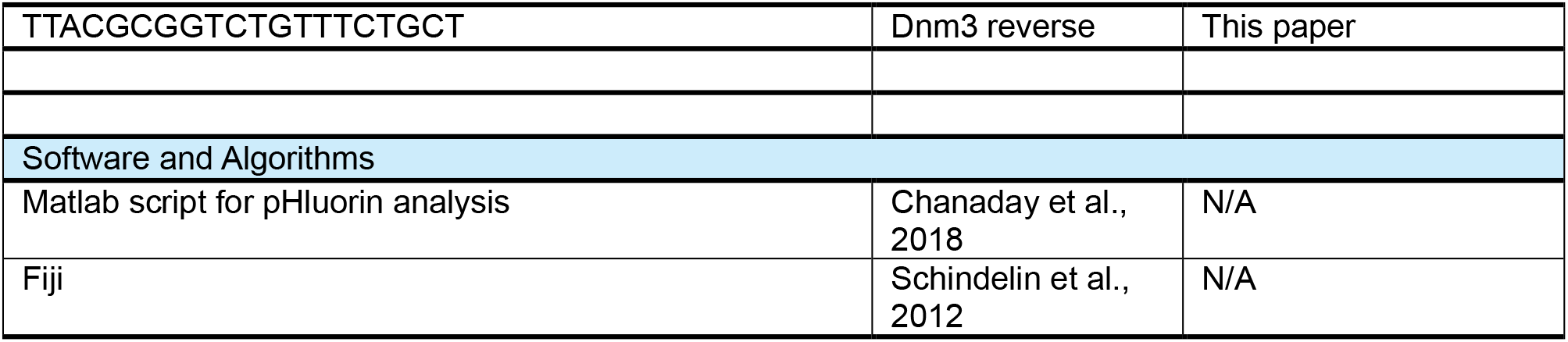

### ETHICS

Animal experimentation: Animal procedures conformed to the Guide for the Care and Use of Laboratory Animals and were approved by the Institutional Animal Care and Use Committee at UT Southwestern Medical Center (Animal Protocol Number APN 2016-101416) and at Vanderbilt University School of Medicine (Animal Protocol Number M1800103).

### METHOD DETAILS

#### Lentiviral infection

HEK293 cells (ATCC) were transfected with 3 lentiviral packaging plasmids (pMDLg/pRRE, pRSV-Rev, and pCMV-VSV-G) and a pFUGW vector containing a Cre-recombinase and/or vGluT1-pHluorin construct using the Fugene 6 transfection reagent (Promega). Cell culture supernatants containing the virus were collected 72 hours later and spun down to precipitate out cellular debris and other contaminants. Neurons were infected at 4 days *in vitro* (DIV) by adding 200 μl of virus containing supernatant to the neuronal culture media.

#### Cell culture

Dissociated hippocampal cultures from postnatal day 0-3 dnm1^f/f^dnm2^f/f^dnm3 KO were prepared as previously described (Kavalali, Klingauf, & Tsien, 1999). Neurons were infected at 4 DIV with lentivirus expressing Cre recombinase or an empty L307 vector for control and experiments were performed at 17-21 DIV. All experiments were performed following protocols approved by the University of Texas Southwestern Institutional Animal Care and Use Committee.

#### Immunocytochemistry

Neuronal cultures were processed for immunocytochemistry as described in Ramirez *et. al.* (2008) at 17-19 DIV. Antibodies against dynamin (Synaptic Systems), dynamin 1 (Abcam), dynamin 2 (Abcam), synapsin, vGlut-1 (Synaptic Systems) and vGAT (Synaptic Systems) were used at concentrations of 1:500, 1:300, 1:300, 1:1000, 1:500 and 1:500 respectively. Images were taken on a confocal microscope with a 60X 1.4 NA objective. 8-10 images were taken from 3 coverslips per group. Colocalization analysis was object based and performed using Fiji software (Schindelin et al. 2012) and a custom made macro.

#### Electron Microscopy

Neuronal cultures were incubated in modified Tyrode’s buffer (145 mM NaCl, 4 mM KCl, 2 mM MgCl_2_, 10 mM glucose, 10 mM HEPES, 2 mM CaCl_2_, pH 7.4, osmolarity 310 mOsM) for 2 minutes and then rinsed twice with PBS and then fixed and processed for electron microscopy by the UTSW Electron Microscopy Core.

#### Western Blot

Neuronal cultures were homogenized processed for western blot as described in Nosyreva and Kavalali (2010). Antibodies against dynamin (Liu et al., 2008), dynamin 1 (Abcam) and dynamin 2 (Abcam) were used at a 1:1000, 1:3000 and 1:800 dilution respectively and protein bands were developed using enhanced chemiluminescence (ECL). Bands were analyzed using ImageJ software and protein levels were normalized to GDI loading control.

#### RNA extraction and quantitative-Reverse Transcriptase (qRT) PCR

To determine relative expression of different dynamin mRNA isoforms after lentivirus infection, we performed qRT-PCR. RNA from the DIV 19 neuronal cultures were collected by using PureLink RNA Mini Kit (Ambion, Cat. # 12183018A). Quality and purity of RNA were confirmed by NanoDrop2000. For cDNA synthesis, we used Invitrogen’s SuperScript™ III Reverse Transcriptase (Cat. #18080-093). Following its first-strand cDNA synthesis protocol, we used Promega’s Oligo(dT)15 (Cat. # C1101) primer and Recombinant RNasin® Ribonuclease Inhibitor (N2511). cDNA concentration and quality were confirmed by NanoDrop2000 before q-RT PCR.

By following Appliedbiosystems PowerUp™ SYBR™ Green Master Mix (Cat. #A25742) protocol, transcripts for Dnm1, Dnm2, Dnm3 and Gapdh were amplified in a Stratagene Mx3005P. Thermal cycling conditions consisted of 1 cycle of 50 °C for 2 m and 95 °C for 2 m, 40 cycles of 95 °C for 15 s, 57 °C for 45s, 72 °C for 60 sec, and 1 dissociation cycle of 95 °C for 15 s, 60 °C for 60 s, 95 °C for 15 s. Respective primers were obtained from Integrated DNA Technologies: 5′-CTACCACAGAATATGCCGAGT-3′ and 5′-ACTGGGACCTTGGTCATTCC-3′ for Dnm1; 5′-TCACACCTGCCAACATGGAC-3′ and 5′-CGCCGATATAGCCTCTTCTCA-3′ for Dnm2; 5′-GAATTTCGTGGGCAGGGACT-3 and 5′-TTACGCGGTCTGTTTCTGCT-3′ for Dnm3; 5′-AGG TCG GTG TGA ACG GAT TTG-3′ and 5′-TGT AGA CCA TGT AGT TGA GGT CA-3′ for Gapdh. We calculated relative quantification by using 2-ΔΔCt. The number 2 in the equation indicates DNA doubling in each cycle.

#### Electrophysiology

A modified Tyrode’s solution was used for all experiments (except where noted otherwise). Pyramidal neurons were whole-cell voltage clamped at −70 mV with borosilicate glass electrodes (3-5 MΩ) filled with a solution containing (in mM): 105 Cs-methanesulphonate, 10 CsCl, 5 NaCl, 10 HEPES, 20 TEA.Cl hydrate, 4 Mg-ATP, 0.3 GTP, 0.6 EGTA, 10 QX-314 (pH 7.3, osmolarity 300 mOsM). Excitatory-postsynaptic currents (EPSCs) were evoked with 0.1 ms, 10 mA pulses delivered via a bipolar platinum electrode in a modified Tyrode’s solution containing Picrotoxin (PTX, 50 μM) and D-2-Amino-5-phosphonovaleric acid (D-APV, 50 μM, NMDA receptor blocker). Spontaneous miniature EPSCs (mEPSCs) were recorded in a modified Tyrode’s solution containing TTX (1 μM), PTX (50 μM) and D-APV (50 μM). Inhibitory-postsynaptic currents (IPSCs) were evoked with 0.1 ms, 10 mA pulses delivered via a bipolar platinum electrode in a modified Tyrode’s solution containing 6-Cyano-7-nitroquinoxaline-2-3-dione (CNQX, 10 μM, AMPA receptor blocker) and D-APV (50 μM). Spontaneous miniature IPSCs (mIPSCs) were recorded in a modified Tyrode’s solution containing TTX (1 μM), CNQX (10 μM) and D-APV (50 μM). Data was analyzed offline with Clampfit 9 software.

#### Imaging

##### vGlut1-pHluorin

17-21 DIV neuronal cultures infected with lentivirus expressing either vGluT1-pHluorin or vGluT1-pHluorin and Cre recombinase were used for imaging experiments. Images were taken with an Andor iXon Ultra 897 High Speed Camera (Andor Technology Ltd) through a Nikon Eclipse TE2000-U Microscope (Nikon) using a 100X Plan Fluor objective (Nikon). Images were illuminated with a Lambda-DG4 (Sutter instruments) and acquired at ~7 Hz with a 120 ms exposure time and binned at 4 by 4 to increase the signal-to-noise ratio. Images were collected and processed using Nikon Elements Ar software prior to export to Microsoft excel for analysis. ROIs were randomly selected based on a threshold after treatment with NH_4_Cl.

##### FM5-95 dye

17-21 DIV hippocampal neuron cultures infected with lentivirus expressing either Cre-recombinase and GFP or an empty L307 vector were used. Cultures were incubated for 2 minutes in a 45 mM K^+^ modified tyrode’s solution containing FM5-95 dye at a concentration of 18μM. Subsequently, excess dye not endocytosed was washed out for 7 minutes with 2 mM Ca^2+^ tyrode’s solution and synaptic vesicle release of dye was stimulated by two perfusions of 90 mM K^+^ modified tyrode’s solution for 90 minutes each separated by a 90 min washing with 2 mM Ca^2+^ tyrode solution. Images were captured with a Nikon Eclipse TE2000-U Microscope (Nikon) using a 40X Fluor objective (Nikon). Images were illuminated with a Lambda-DG4 (Sutter instruments) and acquired at 1 Hz with a 200 ms exposure time. Images were collected and processed using Nikon Elements Ar software prior to export to Microsoft excel for analysis. ROIs were randomly selected.

#### Imaging Analysis

##### vGluT1-pHluorin

Individual synaptic puncta were selected randomly after NH_4_Cl perfusion and all quenched pHluroin probes were unmasked. Single vesicle events were analyzed as reported in Leitz et al. (2011) and Chanaday et al. (2018). Dwell times were calculated as the time between the initial fluorescence step and the start of fluorescence decay (using the first derivative as parameter). Single vesicle events were analyzed offline with Matlab. For 20 Hz stimulation, amplitude measurement and single exponential decay fitting (using Levenberg-Marquardt least sum of squares minimizations) were performed offline in Clampfit.

##### FM dye

Individual puncta were selected randomly after initial washout of excess dye not endocytosed. Synaptic pool size was estimated by taking the difference of the average fluorescence of individual puncta prior to 90 mM K^+^ stimulation from the average fluorescence at the end of the second 90 mM K^+^ stimulation. The rate of FM dye release was calculated by fitting the decay in fluorescence in response to the initial 90 mM K^+^ tyrode stimulation to a single exponential decay function using Levenberg-Marquardt least sum of squares minimizations.

#### Statistical Analysis

Statistical analyses were performed with Graphpad Prism 6 software using one of the following tests: Student’s ordinary t-test, Student’s paired t-test, Two-way RM ANOVA and Kolmogorov-Smirnov test. Error bars represent SEM.

## Acknowledgements

We would like to thank Drs. Helmut Kramer (UT Southwestern) and Sandy Schmid (UT Southwestern) for sharing reagents as well as for their critical insight into this project. This work was supported by National Institute of Mental Health grants, MH081060 and MH070727 (LMM), and MH66198 (ETK).

---

**Figure 1 – figure supplement 1.**
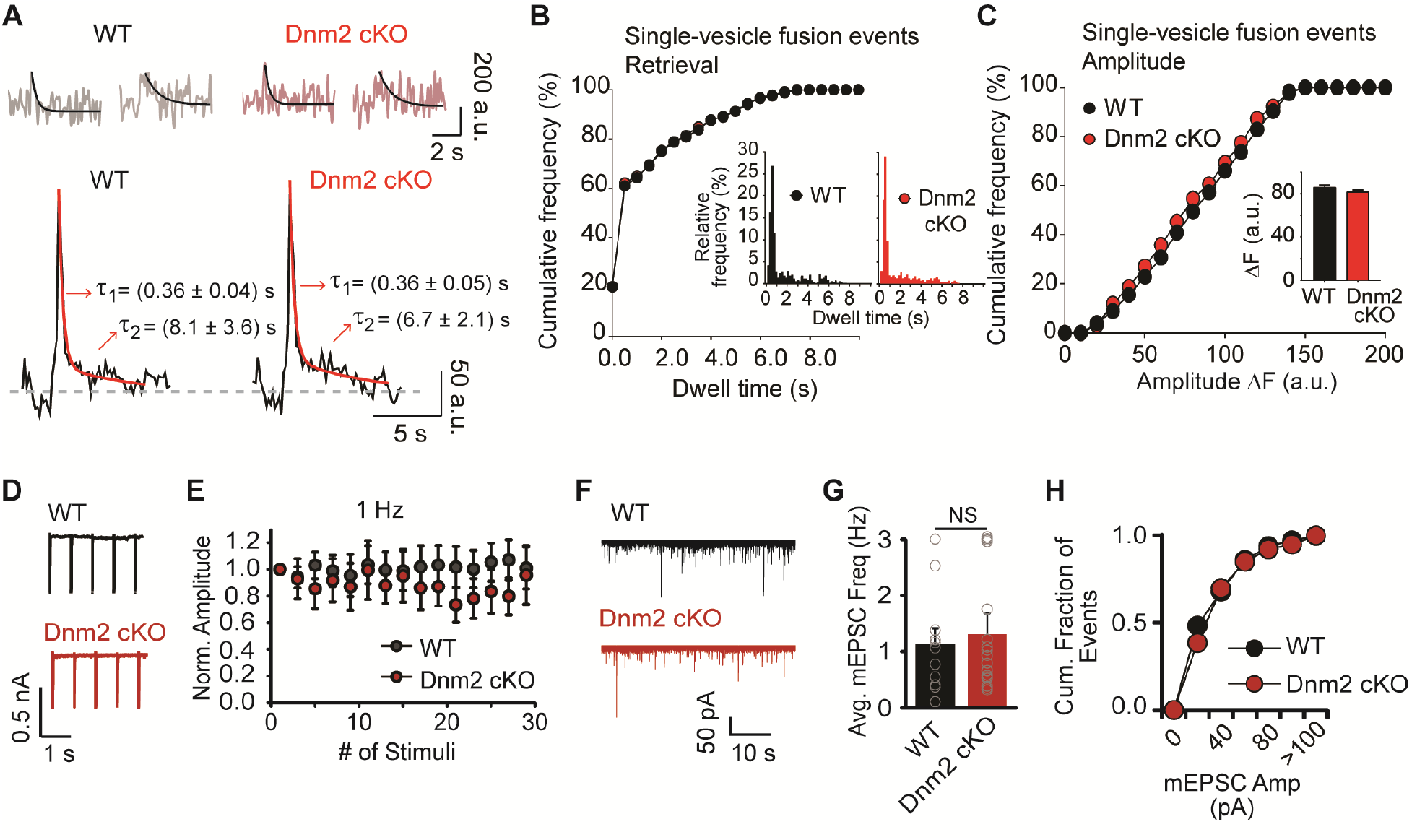
**A.** Analysis of single vesicle fusion events in Dnm 2 KO synapses at extracellular 2 mM Ca^2+^ concentration. Top: two representative single vesicle fusion events monitored with vGluT1-pHluorin for control (WT, grey) and Dnm 2 KO (pink; black lines show the exponential decay of fluorescence after fusion). Bottom: average of all single vesicle event traces for WT (left) and Dnm 2 KO (right) fitted with a double exponential decay (red line) revealing the two components of endocytosis (ultrafast ~360 ms; fast ~6-8 s). There is not a major impact of removal of Dnm 2 in the kinetics of single synaptic vesicle retrieval and reacidification during low frequency stimulation. **B.** Cumulative distribution of single vesicle event dwell times from WT control (black, N=429 boutons) and Dnm 2 KO (red, N=489 boutons) synapses. There is no effect in the kinetics of single synaptic vesicle endocytosis after removal of Dnm 2. Inset: Histograms of dwell time duration for both experimental groups (p=0.1391 by Kolmogorov-Smirnov test). **C.** Cumulative distribution of single vesicle event amplitudes from WT control (black, N=429 boutons) and Dnm 2 KO (red, N=489 boutons) synapses. Inset: Average amplitude for WT (black, mean=86.1 a.u.) and Dnm 2 KO (red, mean=81.7 a.u., p=0.8130 by Kolmogorov-Smirnov test) presynaptic terminals. **D.** Sample traces of the first 5 responses from WT (black) and Dnm 2 KO (red) neurons to 30 stimuli applied at a 1 Hz frequency. **E.** Normalized responses from WT (black, N=10) and Dnm 2 KO (red, N=9) cultured hippocampal neurons after 1 Hz 30 AP stimulation are similar for both groups (p=0.4385, Two-way RM ANOVA). **F.** Representative mEPSC traces from WT (black) and Dnm 2 KO (red) hippocampal neuron cultures. **G.** Average mEPSC frequency for WT (black, N=11, mean=1.1 Hz) and Dnm 2 KO (red, N=15; mean=1.3 Hz) neurons showing no effect of the loss of dynamin 2 on presynaptic spontaneous release rate (p=0.7500, Student’s ordinary t-test). **H.** Cumulative distribution of mEPSC amplitude from WT (black) and Dnm 2 KO (red) neuronal cultures.

**Figure 1 – figure supplement 2.**
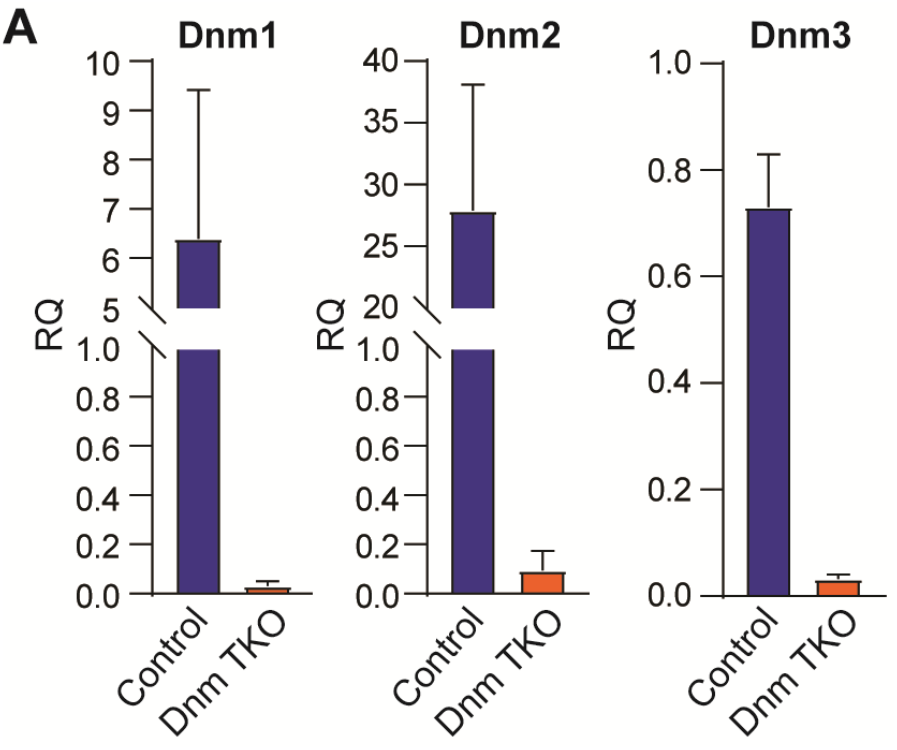
**A.** qRT-PCR of dynamins’ mRNAs in cultured hippocampal neurons from Dnm TKO and littermate controls (controls were floxed Dnm1/2, Dnm3 heterozygous animals incubated with lentivirus expressing only GFP). Data comes from 2 independent cultures at DIV 19, each group was measured by triplicate.

